# Optocoder: computational decoding of spatially indexed bead arrays

**DOI:** 10.1101/2022.02.04.478148

**Authors:** Enes Senel, Nikolaus Rajewsky, Nikos Karaiskos

## Abstract

**Motivation:** Advancing technologies that quantify gene expression in space are transforming contemporary biology research. A class of spatial transcriptomics methods uses barcoded bead arrays that are optically decoded via microscopy and are later matched to sequenced data from the respective libraries. To obtain a detailed representation of the tissue in space, robust and efficient computational pipelines are required to process microscopy images and accurately basecall the bead barcodes.

**Results:** Optocoder is a computational framework that processes microscopy images to decode bead barcodes in space. It efficiently aligns images, detects beads, and corrects for confounding factors of the fluorescence signal, such as crosstalk and phasing. Furthermore, Optocoder employs supervised machine learning to strongly increase the number of matches between optically decoded and sequenced barcodes. We benchmark Optocoder using data from an in-house spatial transcriptomics platform, as well as from Slide-Seq(V2), and we show that it efficiently processes all datasets without modification.

**Availability:** Optocoder is publicly available, open-source and provided as a stand-alone Python package on GitHub: https://github.com/rajewsky-lab/optocoder

## Introduction

Single-cell RNA sequencing methods (scRNA-seq) are by now well-established and of high-throughput, detecting thousands of genes at single-cell resolution^1,2^. Employing scRNA-seq, researchers can readily investigate cellular heterogeneity, cell types and states, and developmental processes for a variety of tissues^3–5^. One shortcoming of all scRNA-seq methods, however, is tissue dissociation that results in loss of spatial context. Spatial information is crucial to study cellular interactions in the native tissue space, to identify spatial expression patterns, and dissect tissue organisation in 3D^6–9^. Such information is essential for the investigation of disease states and progression and it is anticipated that gene expression patterns in space and time will be key for the early detection and interception of complex diseases^10^. In recent years, several efforts have been made to either retrieve the spatial information computationally^11–14^, or to directly sequence gene expression in tissue space experimentally^15^.

One way of acquiring spatially resolved transcriptomics experimentally is to use hybridisation-based methods, such as MERFISH, which achieve single-cell resolution but only for a pre-selected panel of genes^16^ (although this panel can be at genome-scale). In addition to these, sequencing-based techniques that provide unbiased whole-transcriptome spatial data have become available. Methods such as the Spatial Transcriptomics^17^ and the commercially available 10x Visium^18^ use printed spatially barcoded RNA capture probes. In these techniques, however, every spot in space currently aggregates multiple cells. Seq-scope is another method in which Illumina flowcells are used to amplify barcoded oligonucleotides, resulting in a higher resolution system^19^. As a pioneering single-cell resolution platform, Slide-Seq (and Slide-SeqV2), was developed to spatially capture tissue gene expression^20,21^.

In array-based methods, such as Slide-Seq, a tightly packed group of beads carrying DNA oligos are placed on a glass or a plate, termed *puck*. All oligos on the same bead share a random barcode sequence long enough to make this barcode unique for the bead. These barcodes and their positions on the puck are first optically decoded using subsequent rounds of hybridization to fluorescently labelled nucleotides in a microscopy setup^20,21^. After this spatial registration of the beads, a tissue slice is placed on the puck. RNA is captured by the oligos on the beads, amplified, and sequenced-including, for each captured RNA molecule, the bead barcode. Thus, by matching these sequenced barcodes to the optically decoded barcodes, RNA molecules can be mapped to the spatial position of the bead which captured the respective molecules. Similar to Slide-Seq, we are currently also developing a spatial transcriptomics platform using a spatially barcoded assay. Efficient processing and analysis of the acquired datasets takes place in two fronts: in the processing of the sequencing data, for which we have developed Spacemake^22^; and in the processing of the microscopy images.

Computational processing of the microscopy images to retrieve bead barcodes and their locations is challenging and requires three main steps. First, raw images are processed to correct problems such as misalignments across cycles and illumination errors, as well as to detect the beads. Next, the detected beads are processed for basecalling. Several technical issues may distort the signal, such as crosstalk caused by the overlapping laser excitation spectrum and phasing caused by inefficient reactions resulting in lagged signals. Finally, base calling quality is evaluated. Several base calling methods have been developed for sequencing data by primarily modelling the above confounding factors^23,24^. While these methods provide solutions for their respective objectives, there is either no public and easy-to-use implementation, or they are not actively maintained. Hence, there is a lack of a complete pipeline that can process microscopy images from beginning-to-end in an easy-to-use, extensible and robust manner, specifically tailored for array-based spatial transcriptomics assays. In addition, array-based methods require the matching of the optically decoded barcodes to the true set obtained by high-throughput sequencing and the above methods do not make use of such information in a generalizable way.

Here, we developed *Optocoder*, a computational framework to efficiently process microscopy data during the optical sequencing of the barcodes and locations of the arrays in our experimental pipeline. The framework is an open-source Python software package that inputs microscopy images, processes them, corrects confounding factors, such as crosstalk and phasing, and performs basecalling. Importantly, we developed a machine learning based basecaller that increases the number of decoded barcodes that match to sequenced ones. Furthermore, Optocoder employs several measures to control the quality of the decoded barcodes at every processing step. We demonstrate Optocoder’s performance on several datasets, including in-house and publicly available ones, showing the generalizability of the pipeline to different data modalities. Optocoder is scalable, versatile, extendable and can be seamlessly integrated into existing computational pipelines.

## Materials and methods

Optocoder consists of three distinct modules (Fig. 1). The imaging module is used to align the input microscopy images and detect the beads and their respective locations (Fig. 1a). Second, the barcode bases are called by correcting confounding factors, such as spectral crosstalk and phasing (Fig. 1b). Finally, given that a sequencing barcode set is provided, a machine learning classifier is trained to increase the number of barcode matches between the sequencing and the optical set (Fig. 1c). The output of every step is quality controlled with several metrics to create a final report of the puck, image and base calling quality (Sup. Fig. 1 & Sup. Fig. 2).

**Figure 1:**
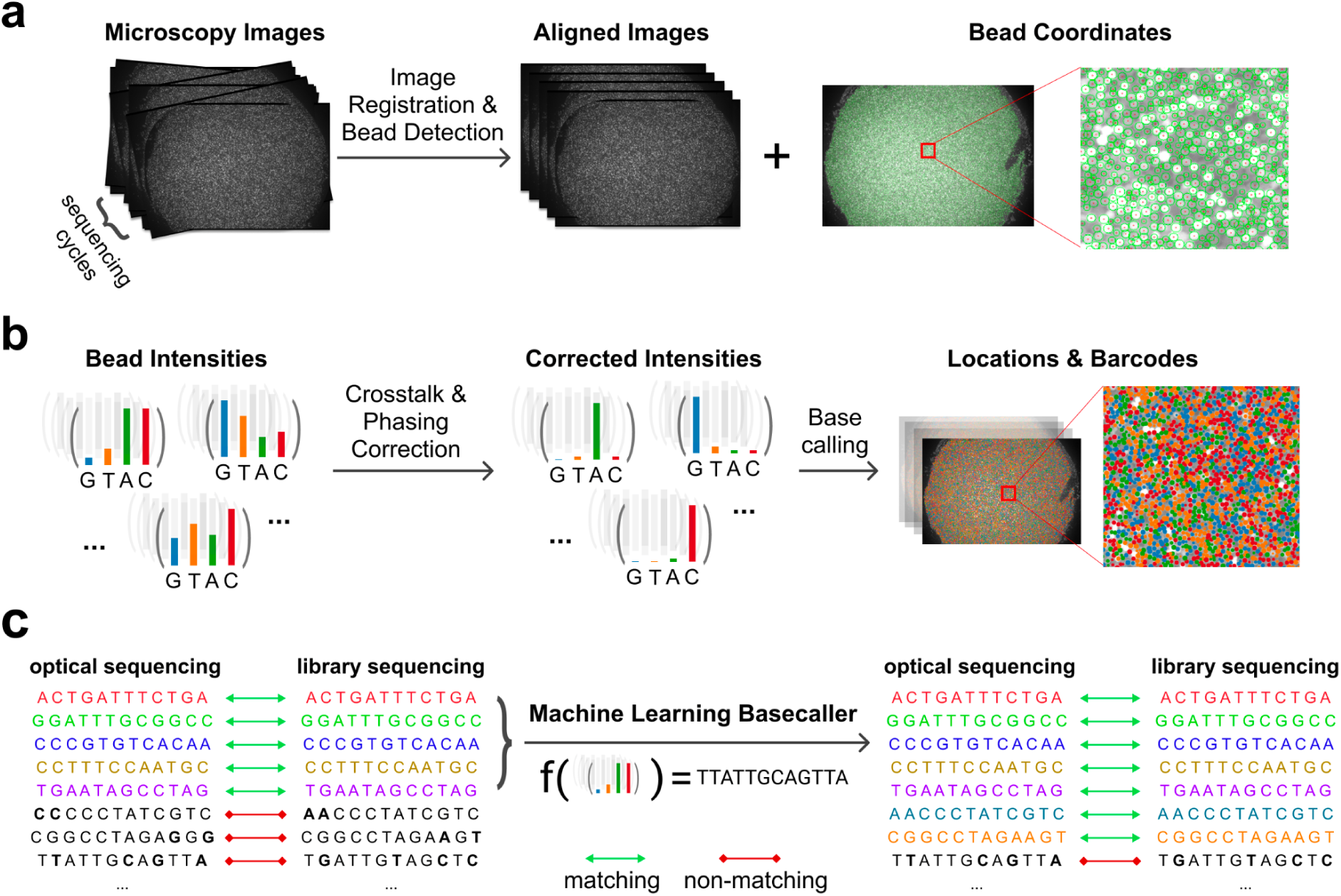
Schematic overview of Optocoder’s modules. **a**, Image processing is used to align microscopy images acquired across the sequencing cycles and to detect the beads and their coordinates on the array. **b**, Crosstalk and phasing effects are corrected for high-quality basecalling. **c**, Machine learning is employed to further correct base calling and increase matches between the optically decoded and sequenced barcodes.

## Image processing

The image processing module is used to align the microscopy images and detect the beads on the array. The input to Optocoder are the puck images acquired via microscopy for every barcode base.

### Bead detection

Beads are adjoining circular-shaped objects of a given radius, and we utilise Hough Circle Transform^25^ to detect them from the overlay image of the last cycle (Sup. Fig. 3a, Methods). For a given bead batch, bead sizes remain constant across experiments, so that Hough Transform requires minimal optimisation. In case a different bead batch contains larger or smaller beads, the image processing module can be readily modified through adjusting the expected bead radius parameter. Bead detection outputs the (x, y) coordinates of the beads which are subsequently used to calculate corresponding channel intensities for each cycle.

### Image alignment

The experimental apparatus can physically move between cycles of optical sequencing, thus resulting in potential positional differences between cycles. To retain the bead identities during the whole sequencing process, the images need to be aligned to be able to assign correct intensities to the detected beads. To begin with, the intensities can vary across cycles and can be very low for the last ones. We therefore first create overlay images and then apply histogram matching for every cycle by using the last cycle as the reference frame. Then, we use an image registration method, Enhanced Cross Correlation Maximization^26^, with a Euclidean motion model to align images from all cycles to a reference and detect warping parameters (Sup. Fig. 3b, Methods). Finally, we evaluate the registration quality for each cycle by using the Structural Similarity Index^27^ (Methods).

### Background Correction

Microscopy images are affected by uneven illumination and background noise that might influence the subsequent image processing and base calling steps. To subtract this uneven background signal, we first detect the background image for every channel separately by using a morphological opening operation and then subtract it from the image (Methods). At the end of the image processing module, Optocoder outputs a matrix containing the 2-dimensional coordinates for each bead on the puck and the average fluorescence intensity for each channel.

## Basecalling

In the absence of technical noise, calling bases could be performed by calling the highest intensity channel’s corresponding nucleotide. As shown in the literature for Illumina sequencing basecallers^23,24,28–30^, however, there exist confounding factors of the microscopy readout that need to be taken into consideration for high accuracy basecalls. Similar issues occur in the case of optical sequencing and we identified spectral crosstalk and phasing effects as the main factors that convolute the signal in our experiments.

### Spectral crosstalk correction

Crosstalk refers to the correlation between the A-C and G-T channels due to overlapping emission spectra of fluorophores excited in two laser microscopy setups (Fig. 2a). Optocoder utilises an estimation method^31^ to detect the overlap between channels (Methods). More specifically, the crosstalk matrix is determined by calculating the intensity overlap between every channel. First, an informative group of bead intensities is selected for every channel which is subsequently fitted with a regression model against the values in every other channel. The slope of these models represent the drift of the intensities towards the other channel and the ratio is subsequently used to correct for crosstalk, resulting in the deconvolution of the A-C and G-T channels (Fig. 2b, Methods).

**Figure 2:**
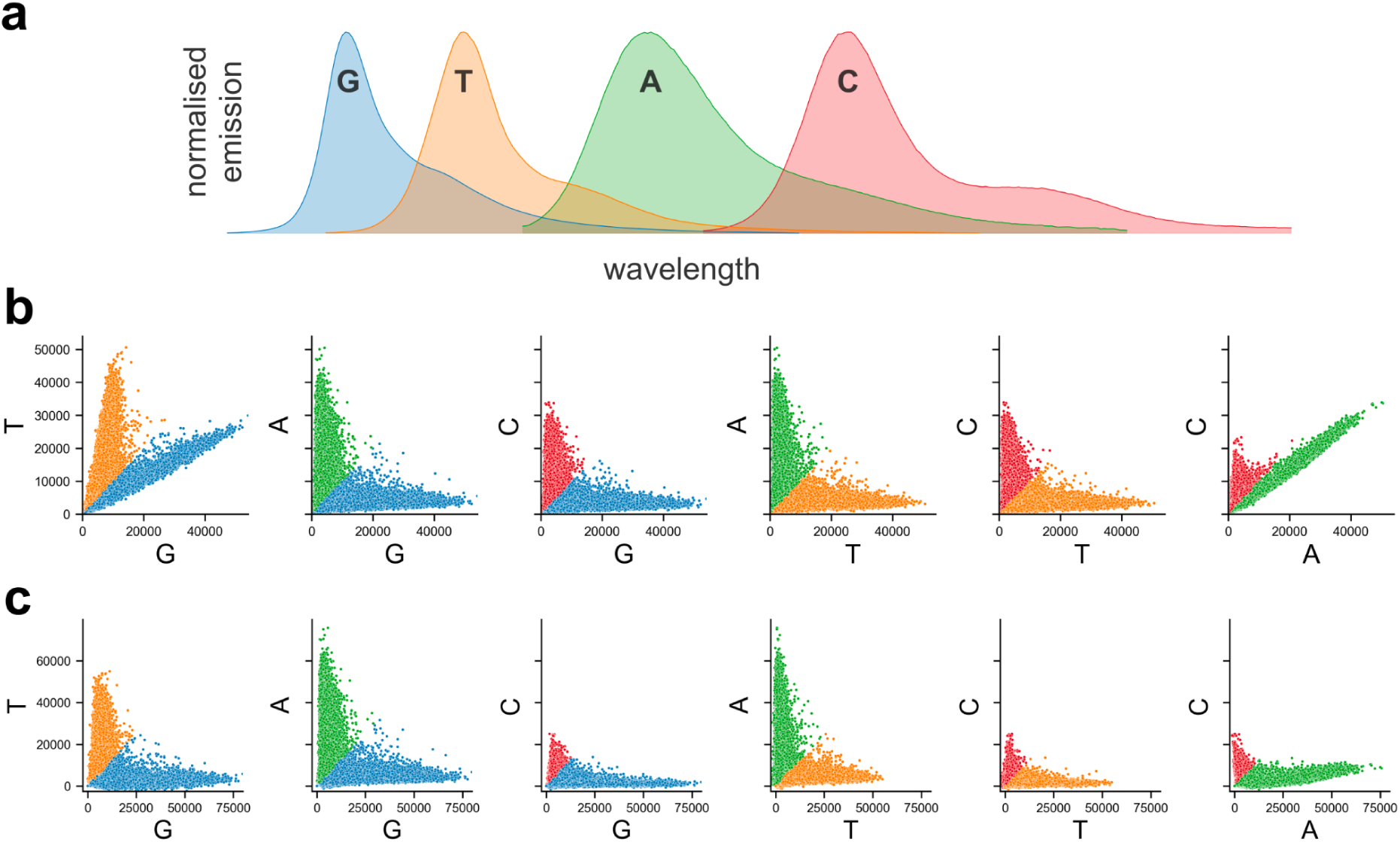
Optocoder efficiently ameliorates spectral crosstalk effects. **a**, The spectra of G-T and A-C channels partially overlap in a two-laser microscopy setup. **b**, Pairwise scatterplots of bead intensities for the puck P4 before (top) and after (bottom) crosstalk correction. Each dot represents a bead and colouring corresponds to the highest intensity of the channel pair plotted for each bead.

### Phasing and prephasing correction

Phasing and prephasing might be caused by inefficient reactions during the nucleotide incorporation process^32^. A bead typically contains millions of oligos that can capture cellular molecules and missing incorporation cycles can take place for several of them. *Phasing* occurs when a nucleotide is incorporated in the next cycle instead of the current one during optical sequencing, so that the signal for that bead lags behind (Fig. 3a-b). Similarly, *prephasing* occurs when multiple incorporations occur within the same cycle and the microscopy readout includes multiple nucleotides at the same time (Fig. 3c).^32^ Phasing and prephasing result in convoluted signals that strongly affect basecalling quality leading to erroneous barcodes sequences. We model such effects through probabilities that correspond to the fraction of bead oligos that have phasing and prephasing for a given cycle (Methods). Subsequently, we construct a matrix that represents the carry over signal among cycles with respect to these probabilities and use it to correct for those effects as described below.

**Figure 3:**
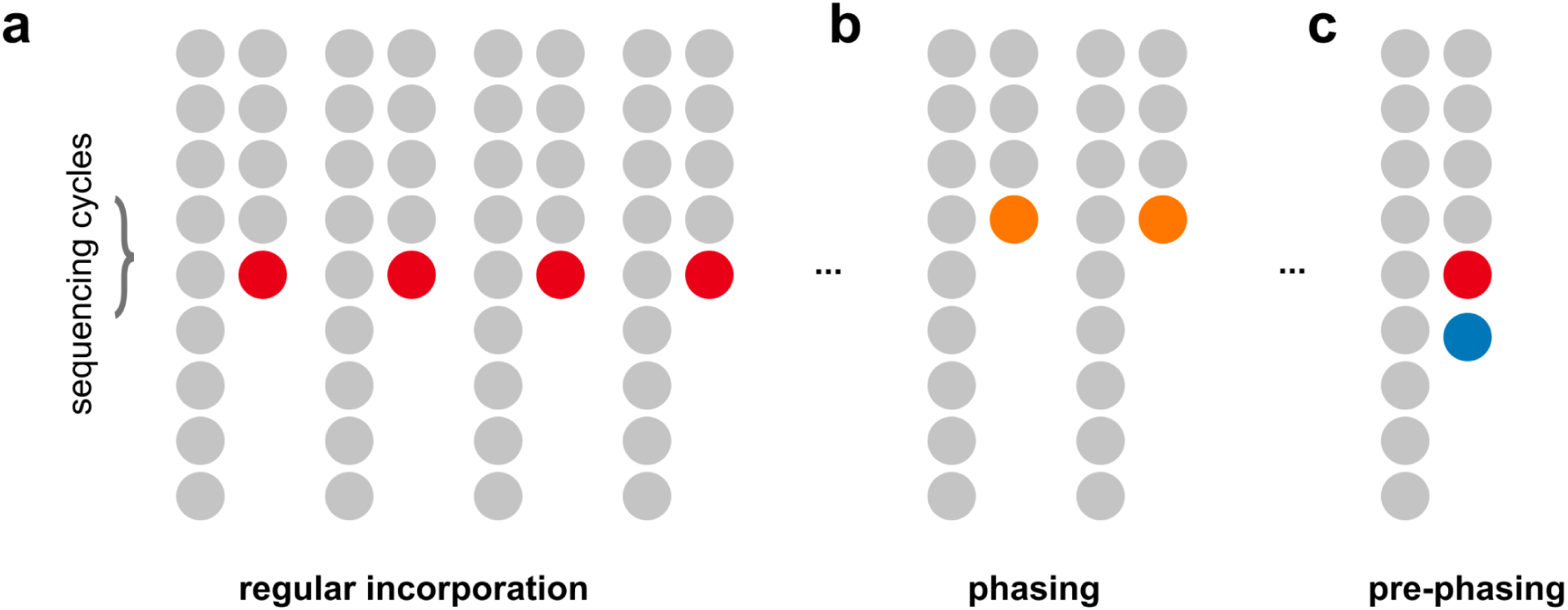
Optocoder efficiently corrects for phasing and prephasing effects. **a**, In the absence of (pre)phasing effects, nucleotide incorporation takes place always in the correct cycle. **b**, Non-incorporation of a nucleotide in the correct cycle results in phasing. **c**, Multiple nucleotide incorporations within the same cycle result in prephasing.

### Combined correction step

To combine the spectral crosstalk and phasing correction, we use a simplified model of the acquired signal similar to^30^ as

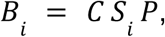

where *B*_*i*_ is a matrix containing the observed intensities of bead *i, C* is the crosstalk matrix, *S*_*i*_ are the true intensities of bead *i*, and *P* is the phasing matrix (Methods). The crosstalk matrix is estimated from the first cycle and assumed to be consistent across cycles since it is a physical phenomenon of the microscopy setup and not cycle dependent. For phasing, Optocoder uses expected phasing and prephasing probabilities chosen by the user. We have observed that for a given bead batch and experimental protocol the amount of phasing and prephasing are consistent across samples (Sup. Fig. 5 & Sup. Fig. 6). However, we have also implemented a phasing parameter search function in which Optocoder determines the best phasing parameters to maximise the number of barcode matches between the optically decoded and the sequenced barcodes. As the intensity ranges of different channels vary, we apply feature scaling for every channel before basecalling (Methods). Optocoder scales channel intensities by removing the median and scaling to the interquartile range (Robust Scaler) and also a normalised exponential function (SoftMax) is applied to each cycle’s intensities for every bead before basecalling (Methods).

### Basecalling and chastity

Having corrected for spectral crosstalk, phasing and prephasing effects, we call barcode bases by selecting the nucleotide of the highest intensity for each cycle. We measure our base calling confidence by computing a chastity score^33^

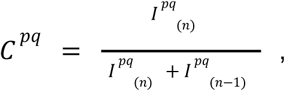

Where *I* ^*pq*^ _(*n*)_and *I* ^*pq*^ _(*n* −1)_ are the intensities of the channels with the highest and thesecond highest values for bead *p* in cycle *q*.

## Machine learning

The spectral crosstalk and phasing corrections greatly improve the basecalling quality and can be readily employed via Optocoder. In array-based spatial transcriptomics methods, however, the true set of barcodes is known via high-throughput sequencing. This provides an opportunity to improve our basecalling by adding a supervised machine learning step. More specifically, we use the optically decoded barcode sequences that exactly match those stemming from the sequencing side as a training set and we train a machine learning classifier for each sample to learn the model parameters that can map bead intensities to nucleotides (Fig. 4). The input features are the background corrected intensities of all cycles after robust scaling for each bead and the model outputs nucleotide probabilities for each cycle and for each bead barcode.

**Figure 4:**
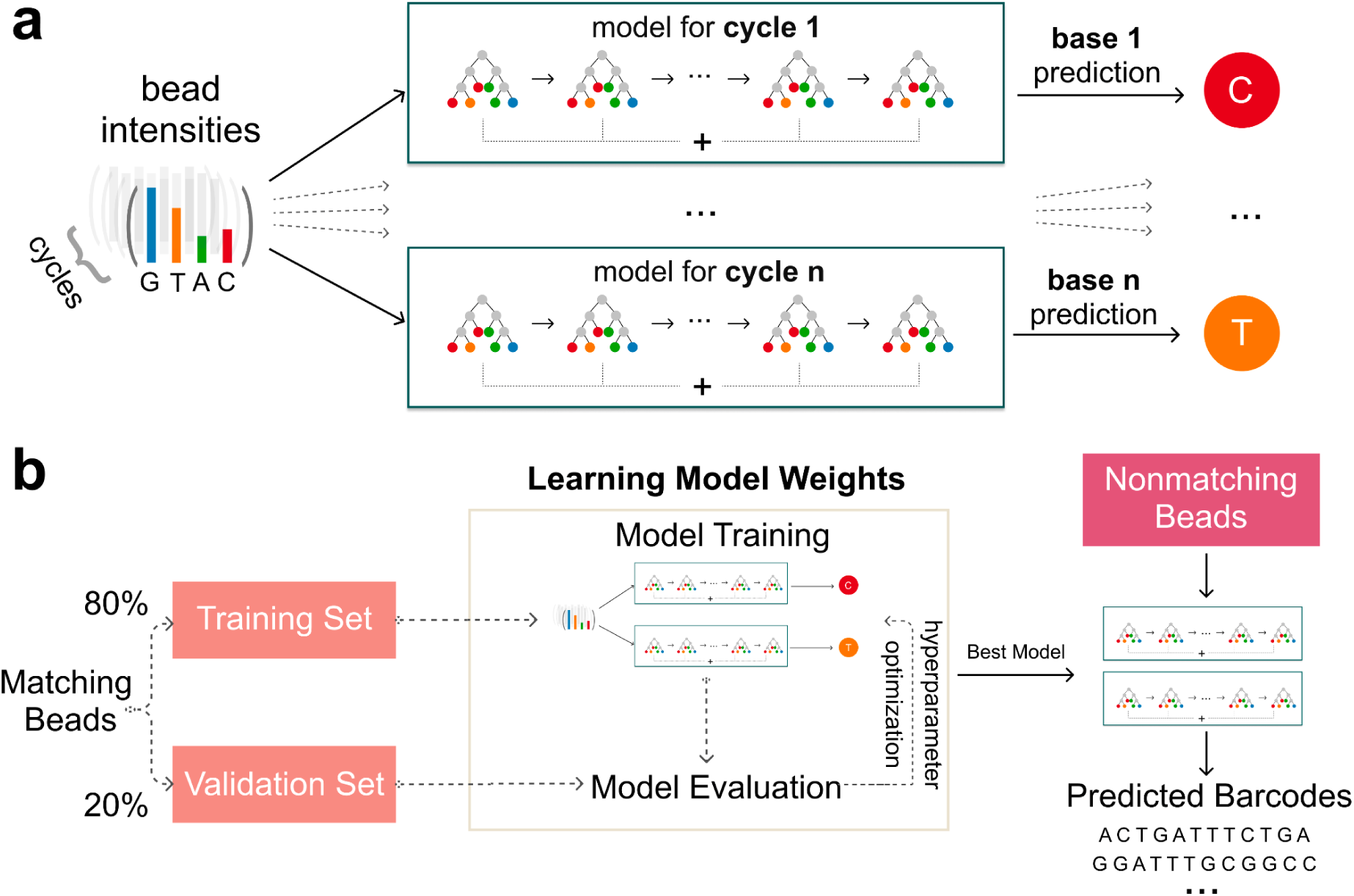
Supervised machine learning increases the number of matches between the optically decoded and the sequenced barcode sets. **a**, a gradient boosting model per imaging cycle is trained to learn and predict nucleotide bases from channel intensities. **b**, Schematic overview of the strategy employed to increase barcode matches.

To efficiently tackle this problem we implemented several classifiers and tested their performance on a number of datasets (Sup. Fig. 4, Methods). We achieved the highest performance by training Gradient Boosting classifiers for each cycle. Gradient Boosting is an additive model that combines weak tree models to improve model accuracy. As we input all cycle intensities to each model, models capture the effects of other cycles’s intensities as well.

We begin with splitting the matching barcodes set into a randomised training (80%) and validation (20%) set. The training set is used as an input into the multi-output Gradient Boosting classifier and the validation set is used to evaluate the model’s performance for hyperparameter optimization. The model with highest accuracy is then retained and used to predict nucleotide bases in the set of non-matching barcodes, which is practically the test set. We evaluate the performance in the test set by computing the number of additional matches to the sequenced barcodes.

### Quality control (QC)

We have implemented several quality controls in Optocoder. These are collectively shown in an automatically generated QC sheet that is associated with each sample (Sup. Fig. 1 & Sup. Fig. 2). In particular, the QC sheet starts with a plot of the raw channel intensities per cycle (Sup. Fig. 1a), which facilitates experimental troubleshooting in case significant cycle deviations occur. Next, the registration accuracy score is plotted (Sup. Fig. 1b) to inspect the quality of the acquired microscopy images per cycle.

After basecalling, the overall nucleotide distribution averaged over all barcodes is plotted to measure the base content (Sup. Fig. 1c). To ensure that the barcode sequences obtained after basecalling are meaningful, Optocoder calculates two measures: string compression and Shannon entropy (Methods). The distributions of these measures are then plotted in the QC sheet against the theoretical distributions expected for randomly uniformed sequences (Sup. Fig. 1d,e). Large deviations from the theoretical distributions would flag low confidence barcode sequences. Additionally, Optocoder plots these measures across puck space, so that areas with low confidence bead barcodes may be identified to evaluate if there are any location-specific barcode quality issues (Sup. Fig. 2c,d).

Furthermore, the distributions of chastity scores that reflect Optocoder’s confidence on basecalling are plotted (Sup. Fig. 1f). Optocoder visualises the distributions of some of these metrics in the array space as the spatial distribution might facilitate a better understanding of the current experiment and also better troubleshooting. Mainly, called bases and their spatial distribution (Sup. Fig. 2a), and the respective chastity scores (Sup. Fig. 2b) for each cycle are plotted and saved.

## Results

### Performance on our data

Our array-based experimental protocol shares certain similarities with SlideSeqV2^21^ and generates microscopy images containing 4 channels for every imaging cycle. These channels display specific fluorescence profiles used to call nucleotides - 4 channels for the four bases G, T, A and C. After library preparation and high-throughput sequencing, the true set of bead barcodes becomes available, so that we can assess Optocoder’s performance.

We benchmarked Optocoder on 4 different pucks that were prepared according to our protocol. Each puck contained around 70,000 beads labelled by 12 bases long barcodes and was optically sequenced under the same experimental conditions. After optical sequencing the following material was placed on the pucks: ERCC spike-ins (P1) or sections of E12 mouse brain (P2, P3 and P4). The prepared libraries were sequenced on an Illumina NextSeq 500 machine and analysed with spacemake^22^ (Methods). The 100,000 beads with the most sequencing reads were considered for matching.

Naive basecalling, i.e., without correcting for crosstalk or phasing, resulted in a low number of matches for all four pucks (Fig. 5a, orange bars). Correcting for spectral crosstalk increased the number of matches by 10% to 20% (Fig. 5a, purple bars). As expected, this increase was reflected by the corresponding chastity scores (Fig. 5b). After crosstalk correction, Optocoder corrected for phasing and prephasing. This step resulted in additional matches for all four pucks. (Fig. 5a, green bars) and a corresponding increase in the chastity scores (Fig. 5b). We observed little to no pre-phasing in our datasets, but a strong phasing effect (Sup. Fig. 5). The combined correction step enhanced the total number of matches by 2-fold compared to naive basecalling.

**Figure 5:**
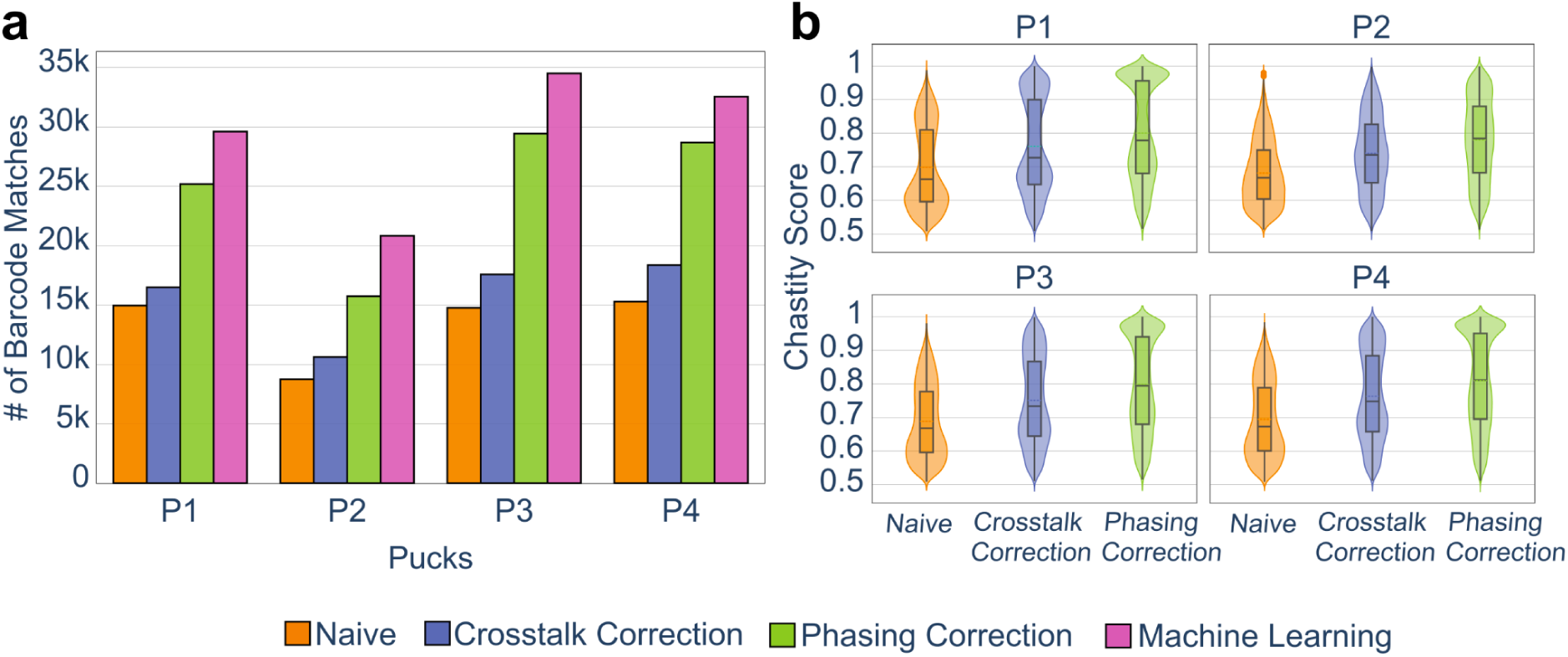
Optocoder exhibits high performance on own-generated data. **a**, Optocoder efficiently corrects for crosstalk and phasing effects, and employs machine learning, resulting in a stark increase of the number of matched barcode sequences compared to naive basecalling in four different pucks. **b**, Chastity scores consistently increase after correcting for crosstalk and phasing effects across all four pucks.

Finally, we trained machine learning models to further increase the number of matches. A model was trained for each puck separately and was used to predict the nucleotides of the non-matching barcode sequences. Optocoder’s machine learning step resulted in a further 15%-32% increase of matches to the combined correction matches (Fig. 5a, pink bars). Interestingly, machine learning performance varied across the four pucks, with the highest increase in number of matches taking place for P2, the puck with the fewest overall matches.

In summary, Optocoder successfully corrected for crosstalk and phasing effects in our datasets and strongly enhanced the number of matches between the decoded and the sequenced barcodes.

## Performance on external data

We primarily developed Optocoder for efficiently processing spatial transcriptomics data stemming from our in-house experimental method. Optocoder, however, is built to be versatile and adaptable. To showcase its flexibility, we used Optocoder to analyse similar microscopy datasets that are publicly available.

The initial Slide-Seq protocol^20^ uses SOLiD chemistry. The optical sequencing images are generated via 20 ligations where 6 of them are constant bases. We applied Optocoder to process the microscopy images associated with three pucks SSP1, SSP2, SSP3 (Sup. Table 2). After image processing, Optocoder detected and identified 52,754 / 48,564 / 63,643 beads, respectively. We observed little-to-no phasing and prephasing effects in the three pucks (Sup. Fig. 6). We compared the decoded barcodes against the true set of sequences that we extracted from the associated BAM file (Methods). Matching the two barcode sets for SSP1 after crosstalk and phasing correction resulted in 31,308 exact matches which is ∼%37 higher than the number of exact matches with Puckcaller, the computational pipeline that was developed and accompanied the Slide-Seq protocol. For SSP2 and SSP3, Optocoder performed similarly to Puckcaller barcodes, with -7% and +5% difference, respectively. By training the corresponding machine learning model, Optocoder resulted in 7% to 58% more matches compared to the baseline Puckcaller basecalling. (Fig. 6).

**Figure 6:**
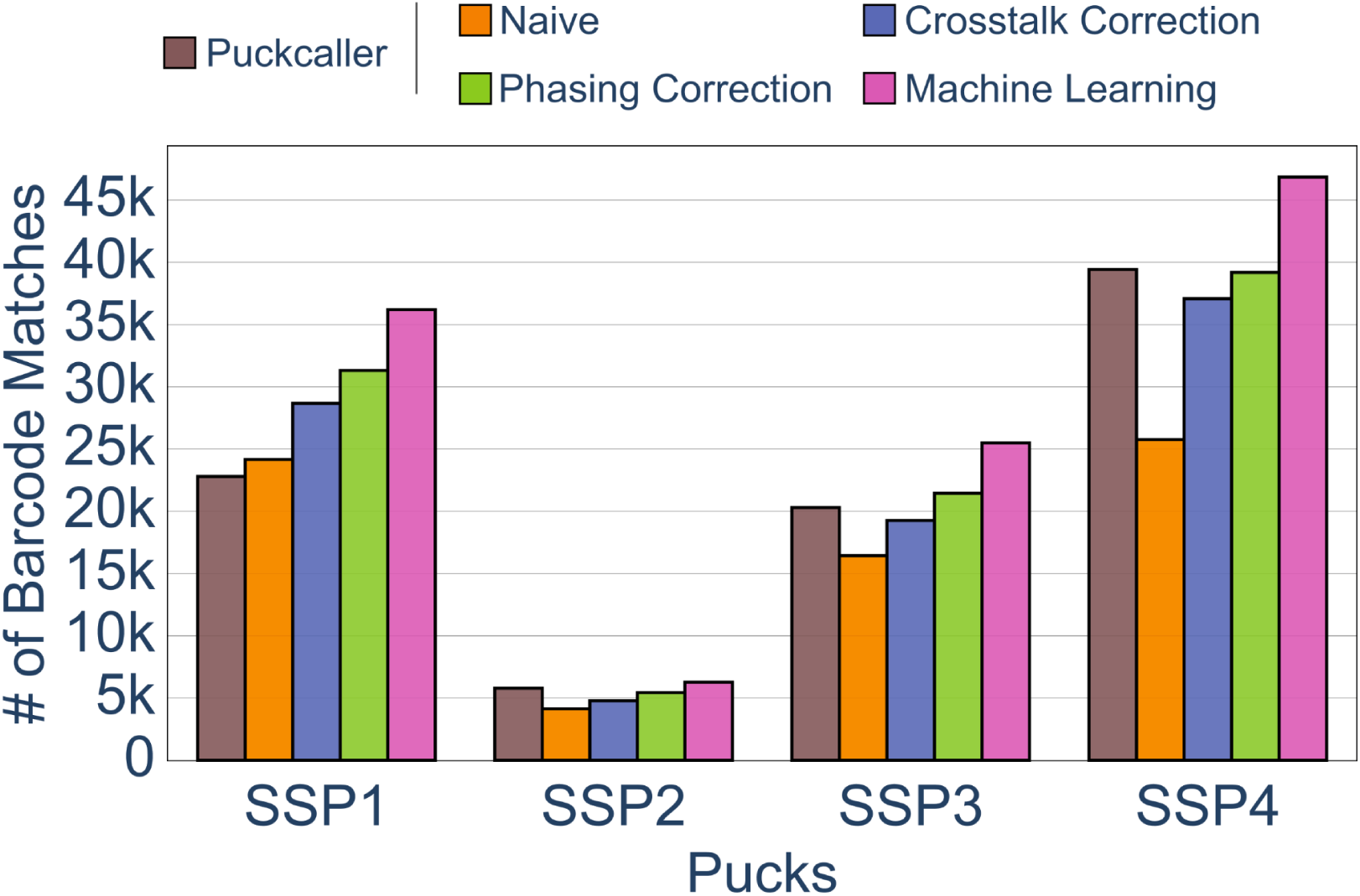
Optocoder efficiently processes published Slide-Seq and Slide-SeqV2 datasets. The number of matches between optically decoded and Illumina sequenced barcodes is shown for three Slide-Seq and one Slide-SeqV2 puck. In all cases Optocoder outperforms Puckcaller, the script used by the authors in the original publications.

In addition to above, we employed Optocoder to analyse Slide-SeqV2 microscopy datasets. This is a sequence-by-synthesis generated microscopy data and the cellular barcodes consist of 14 nucleotides. Optocoder readily processed the microscopy images and decoded 63,643 barcodes for the puck SSP4 (Sup. Table 2). Little-to-no phasing and prephasing effects were also observed for that puck too (Sup. Fig. 6). Comparing the decoded barcodes against the true set of sequenced barcodes that we obtained from the associated BAM file (Methods) resulted in 39,188 exact matches, similar to what the authors acquired for the same dataset. Furthermore, training the machine learning model starkly enhanced the number of matches by ∼17%, resulting in a total of 46,329 exact matches.

Taken together, the above demonstrates that Optocoder can reliably analyse different types of datasets, such as microscopy data of different chemistry, and achieve higher performance than existing methods.

## Discussion

Spatial transcriptomics methods such as Slide-Seq use spatially barcoded bead arrays that are optically sequenced. We anticipate an increase in both the development of similar methods and the utilisation of these methods in various research labs for biological insights. In this study, we present Optocoder, a computational pipeline that efficiently processes microscopy images for optical sequencing of bead barcodes. Optocoder is an easy-to-use, open-source Python package and provides a complete pipeline that processes raw microscopy images to assign bead barcodes in space. Optocoder provides functions to align images, detect beads, correct crosstalk and phasing issues and finally call the bases. Furthermore, we implemented a machine learning pipeline to increase the number of barcode matches between the optically decoded and library sequencing barcodes. We have implemented and compared four different models that are trained separately for each sample and we show that the machine learning approach substantially increases the number of matches.

We initially developed Optocoder for our in-house spatial transcriptomics platform and we evaluated Optocoder on four different samples. Additionally, we have tested Optocoder performance on Slide-Seq and Slide-SeqV2 samples and demonstrated that Optocoder efficiently processes different datasets and experimental setups with minimal modifications. In particular, we showed that correcting for crosstalk and phasing effects improved basecalling quality for our datasets, whereas for Slide-Seq datasets Optocoder performed similarly to what was originally reported. Employing Optocoder’s machine Learning module, however, resulted in a stark increase of barcode matches for both in-house and Slide-Seq datasets.

One drawback of the current crosstalk and phasing correction pipeline is that it implements a linear model and also the correction parameters are not tailored to beads. Beads with unique phasing properties would therefore not be efficiently processed with this approach. As a future development, implementing a more complex model that would allow for bead specific correction parameters might be beneficial.

Improved performance with the machine learning models indicates that nonlinear interactions are not fully captured by the crosstalk and phasing correction model. For machine learning basecalling, one model for each sample is trained to provide a sample-specific basecalling tool without the need for a general training set. However, this approach requires samples that already have a high number of matches before machine learning, so that a model can be trained accurately. For example, relatively poor machine learning performance for SSP2 might be explained by the small size of the initial matching set. While our current model provides a general pipeline that can be used for any new dataset and platform, an additional general model that can be trained commonly and used for different samples might be beneficial. Investigating what is learned by the machine learning models can prove useful to analyse and troubleshoot the experimental reasons for basecalling errors.

Finally, machine learning tools have been previously used for basecalling from raw signals in different platforms such as Illumina and Nanopore to improve basecalling quality ^29,34,35^. In principle, the machine learning approach described here utilises matched sequences and can be potentially extended to such platforms to further improve the basecalling quality by using already called reads in specific contexts, such as genome mappability.

## Author contributions

N.K. conceived and together with N.R. and E.S. designed the initial version of the pipeline. E.S. implemented and developed the pipeline. N.K. conceived and together with E.S. designed the machine learning module, which was then implemented by E.S. E.S. performed all computational and data analyses apart from the processing and analysis of Illumina sequenced libraries that was performed by N.K. N.K. and N.R. supervised the study. All authors wrote the manuscript.

## Code availability

Optocoder is publicly available, open-source and provided as a stand-alone software package on GitHub: https://github.com/rajewsky-lab/optocoder.

## Data availability

We have deposited the microscopy images for P1-P4 on Zenodo under the DOI 10.5281/zenodo.5850814, and the corresponding Illumina sequencing data on GEO under the accession number GSE193472. The Slide-Seq microscopy images and sequencing data were downloaded from the Single Cell Portal. The Slide-SeqV2 microscopy images were provided by Evan Murray and Evan Macosko and the sequencing data were downloaded from the Single Cell Portal.

## Acknowledgements

We are indebted to our colleagues S. Abbiati, J. Alles, S. Ayoub, A. Boltengagen, A. Diag, S. Ehrig, M. Jens, J. Licha, G. Macino, M. Schott, T.R. Sztanka-Toth, S. Tagliaferro and A. Woehler for the adaptation and further development of the Slide-seq protocol, as well as the whole Rajewsky Lab for discussions. We would also like to thank Evan Murray and Evan Macosko for kindly providing the Slide-SeqV2 microscopy images. N.K. was supported by the DFG grants KA 5006/1-1, RA 838/5-1. E.S. was supported by the DFG grant KA 5006/1-1. All authors were supported by MDC Berlin.

