## Supplementary Figures for "Optocoder: computational decoding of spatially indexed bead arrays"

**Sup. Fig. 1**: Example of QC Plots shown for P4. **a,** Raw channel intensities across cycles. **b,** Similarity scores for registered and unregistered cycle images. Cycle 12 exhibits a perfect score, as previous cycles are registered to that. The dashed line at 0.5 denotes an empirical threshold. **c,** Called base fractions per cycle. **d,** Compression score distribution of the barcodes along with the expected distribution. **e,** Entropy score distribution of the barcodes along with the expected distribution. **f,** Basecalling chastity score distribution per cycle. **g,** Histogram of the beads that has n cycles higher than the threshold. **h,** Histogram of the base positions where a base is called with a score lower (red) or higher (blue) than the threshold. **i,** Scores per cycle for two randomly chosen beads.

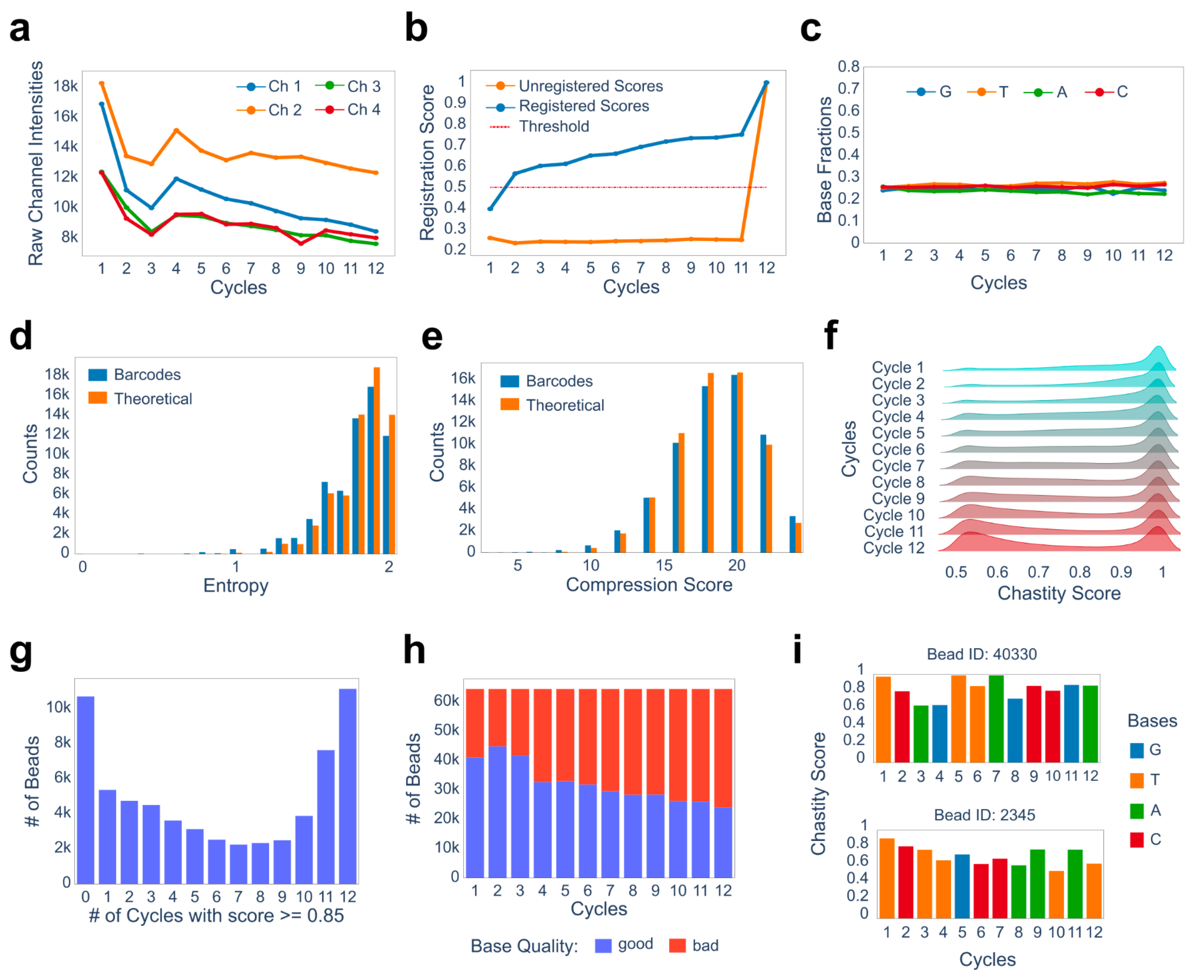

**Sup. Fig. 2**: Example of QC Sheet Plots shown for puck P4. **a,** Called bases in space for every cycle. Here shown cycles 1, 6 and 12. **b,** Chastity scores in space for every cycle. Here shown cycles 1, 6 and 12. **c,** Entropy score distribution in space. Higher values correspond to higher barcode complexity. **d,** Compression score distribution in space. Higher values correspond to higher barcode complexity.

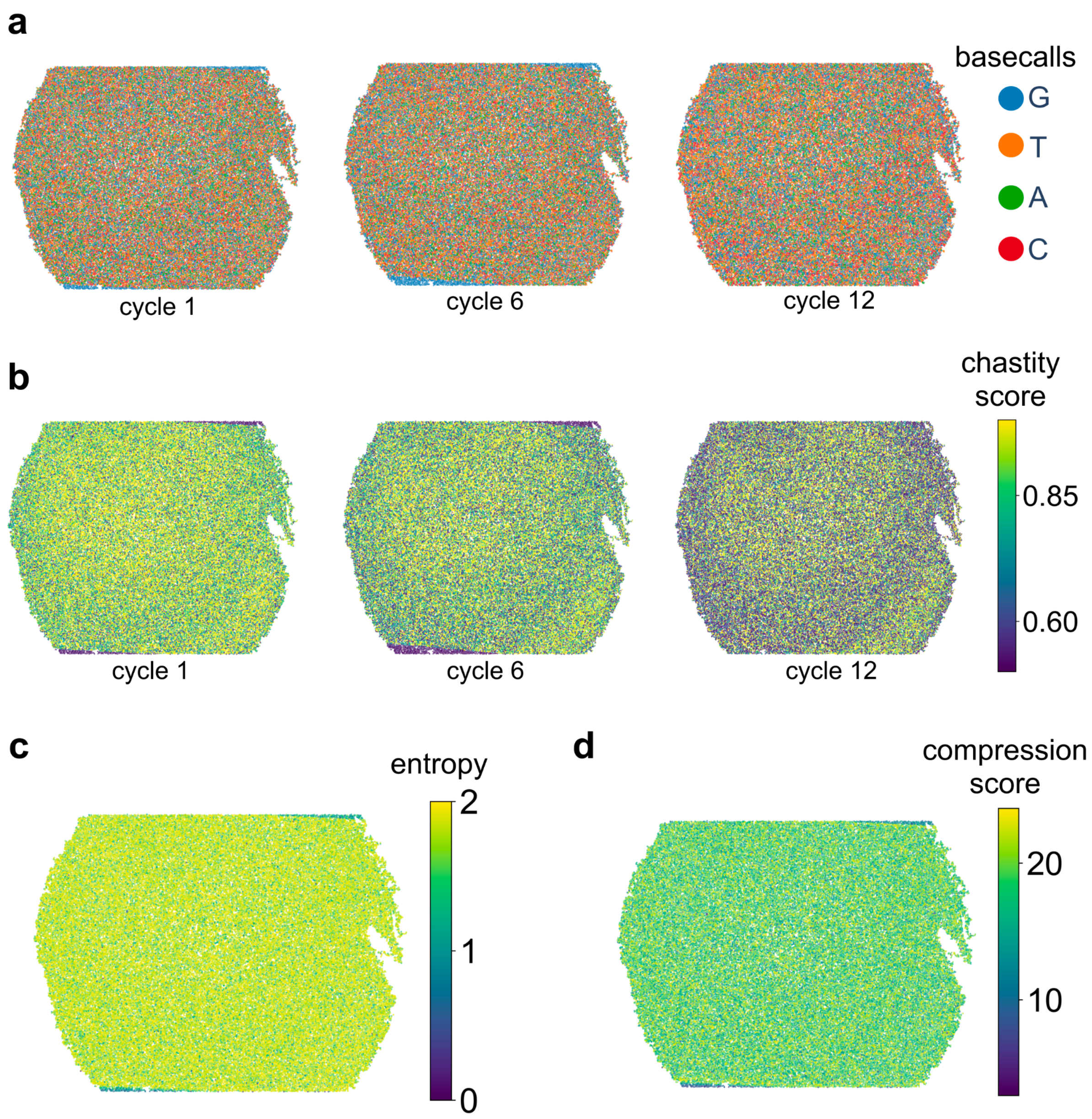

**Sup. Fig. 3:** Registration and bead detection examples shown for P4. **a,** Beads are detected from the reference image which is the overlay of all channel intensities of the last cycle. **b,** Histogram matching and image registration examples shown for cycles 1, 6, 11 and 12.

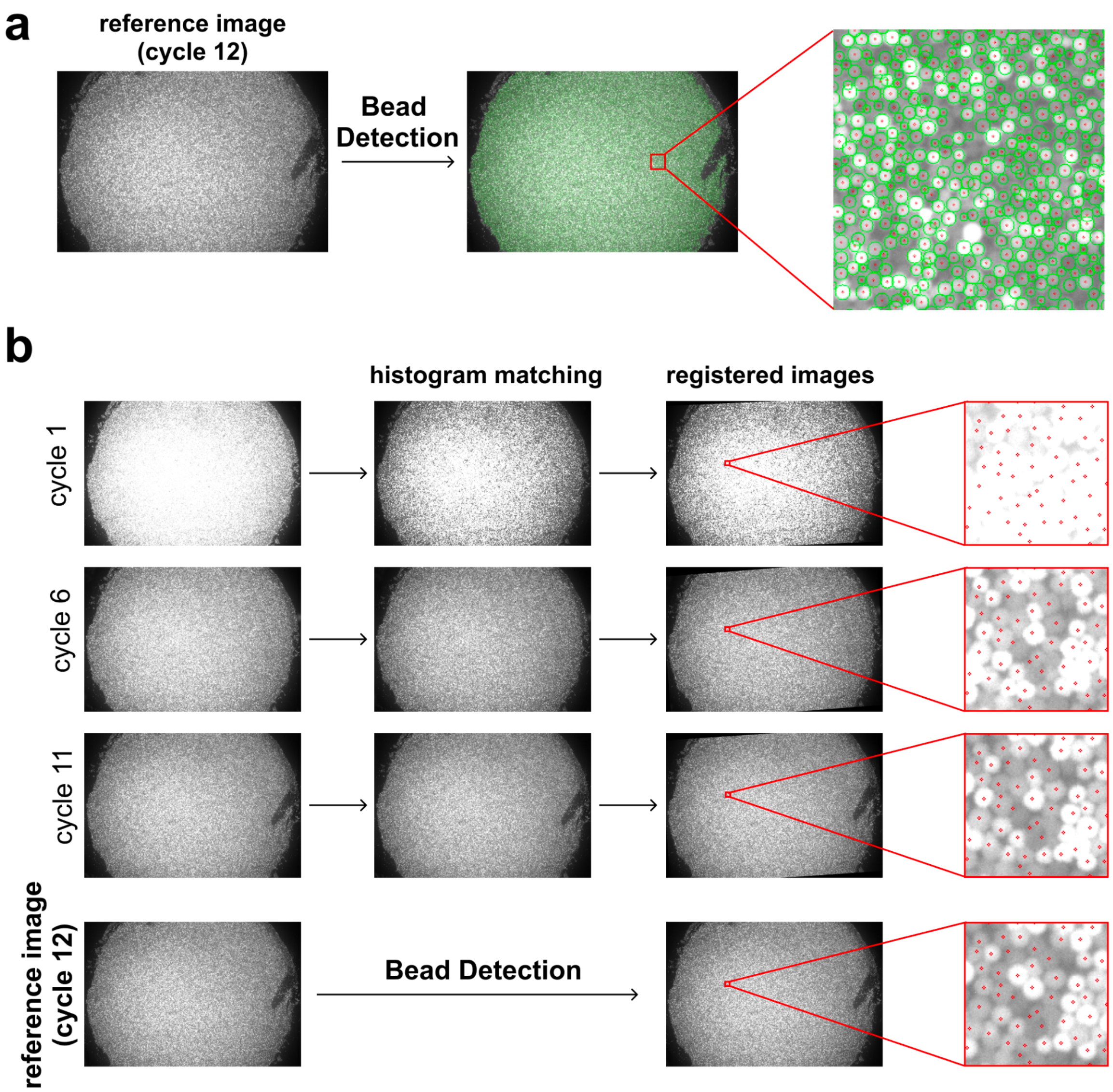

**Sup. Fig. 4:** Benchmarking the performance of various classifiers for the machine learning basecaller. **a,** Number of matches between the optically decoded and the Illumina sequenced barcodes for the in-house pucks and for four different machine learning classifiers. **b,** Same as in a, but for the Slide-Seq and Slide-SeqV2 pucks.

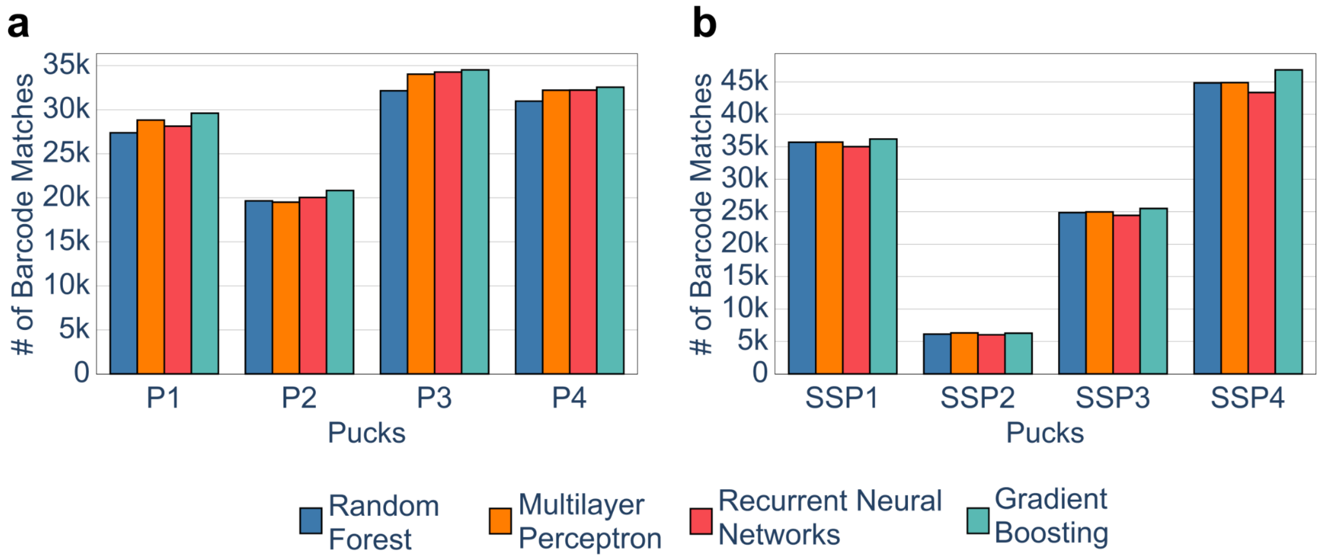

**Sup. Fig. 5:** Heatmaps showing the number of matches between the optically decoded and Illumina sequenced barcodes as a function of phasing and prephasing probabilities for the pucks P1-P4. Moderate phasing and no pre-phasing effects are observed across all pucks.

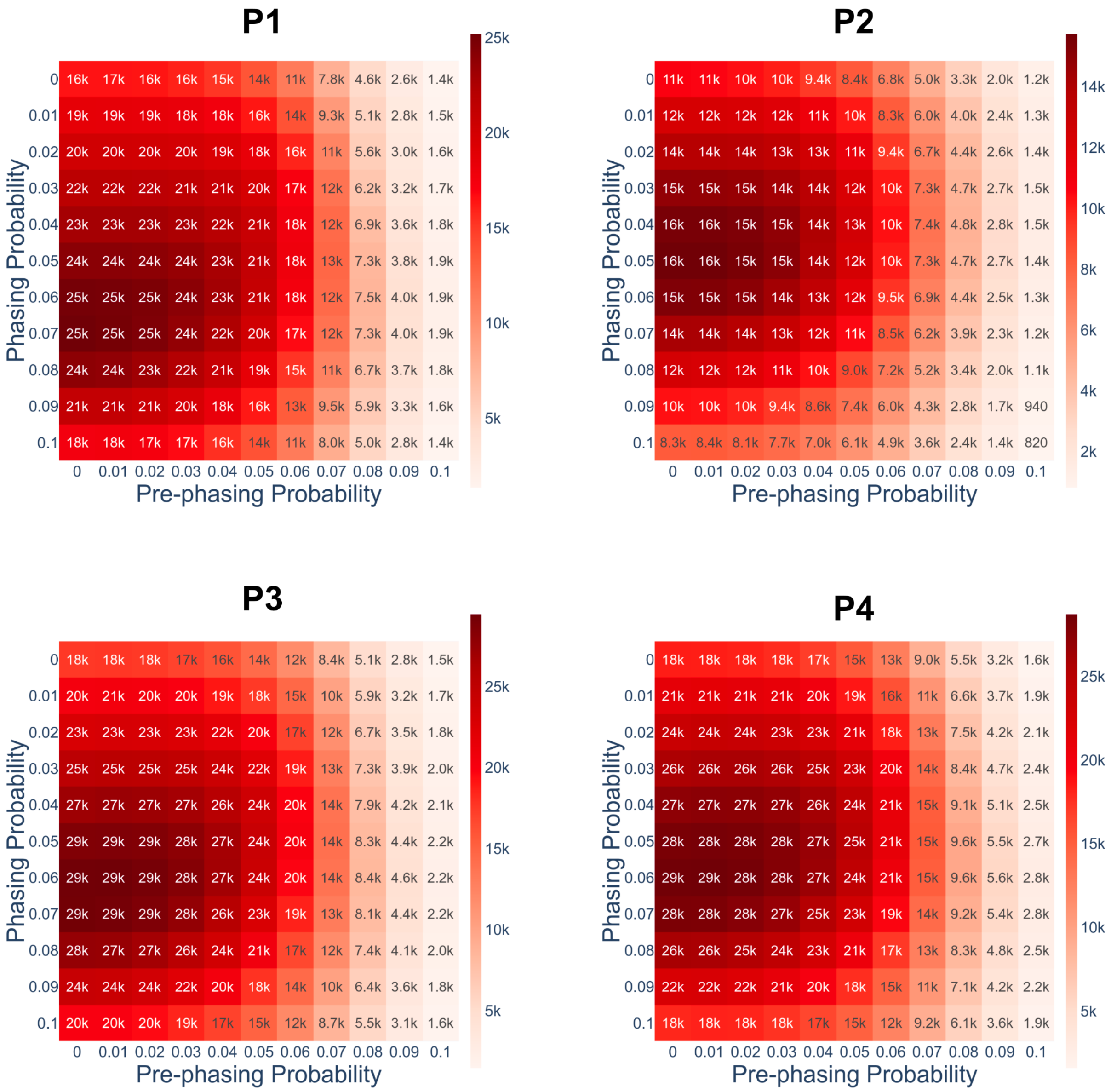

**Sup. Fig. 6:** Heatmaps showing the number of matches between the optically decoded and Illumina sequenced barcodes as a function of phasing and prephasing probabilities for the Slide-Seq and Slide-SeqV2 pucks. Little-to-no phasing and pre-phasing effects are observed across all pucks.

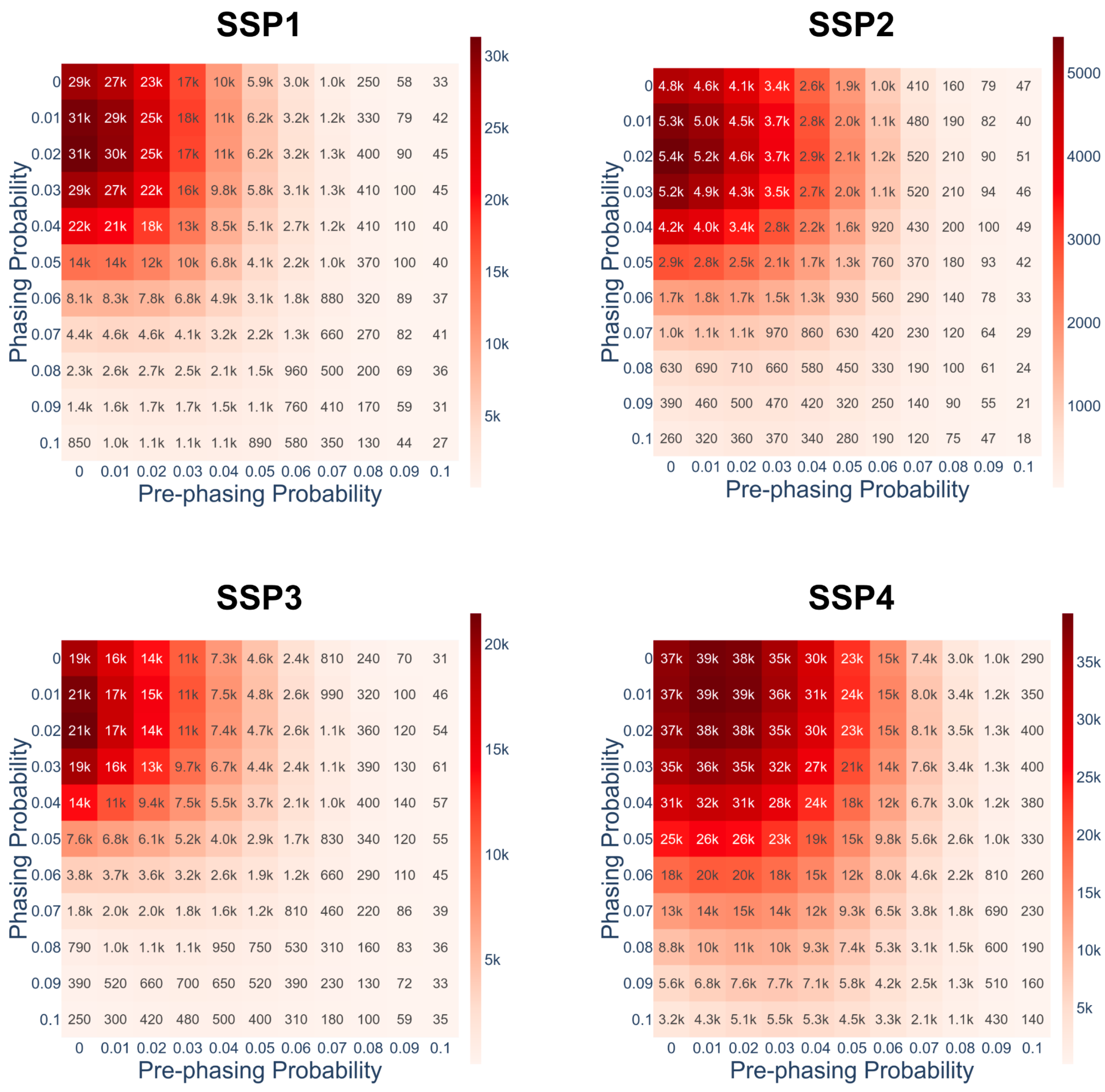
