## Supplementary Methods for "Optocoder: computational decoding of spatially indexed bead arrays"

### Image Processing

Puck images generated through the in-house platform consist of 6-channel images for every cycle, where 4 channels were retained for further processing and the other two were ignored since they were only used for internal microscopy setup control purposes. Pucks from Slide-Seq and Slide-SeqV2 datasets are 4-channel images which were directly used for further processing. For Slide-Seq, a TIFF file is provided per channel and we combined the channels to create a multipage TIFF image for each cycle. For Slide-SeqV2 raw microscopy data we used Puckcaller (https://github.com/MacoskoLab/PuckCaller) to generate the stitched TIFF files. After loading the images, an overlay image for each cycle was generated by summing up the intensities of all channels and was subsequently used for bead detection and image registration. Image processing operations described below were implemented with the OpenCV (v4.2.0) and Skimage (v0.18.3) packages.

#### Bead Detection

Beads were detected from the last cycle's overlay image. First, a median blur (3x3) for noise removal and contrast limited adaptive histogram equalisation (CLAHE) with 64x64 tiles were applied to the overlay image. Then, Optocoder used the Hough Circle Transform OpenCV implementation to detect beads. By default, Optocoder searches for beads with a radius range of 4-10 pixels. These parameter and edge detection thresholds for Hough Transform can be modified through the bead detection parameters as necessary. For the results presented in this paper, we used the same parameters in all runs for both in-house and Slide-Seq datasets. Furthermore, puck borders were detected by searching for the biggest connected component in the image after applying a Gaussian blur. Detected beads outside of the main puck area were ignored.

#### Image Registration

Every image was registered to the final cycle to correct misalignments between cycles. First, a histogram matching step was applied. This was done via calculating the cumulative distribution function for both the image-to-be-registered and the reference and consecutively matching them. Image registration was done via the Enhanced Cross Correlation Maximization[1] algorithm implementation in OpenCV with a Euclidean motion model. Registration was done with a pyramidal scheme where the image was downsampled eight times to speed up the registration process and improve the accuracy by aligning the image in different resolutions, starting from the coarse to the finer resolutions. Warping parameters were acquired from the registration of the overlay images and they were consecutively applied to the channel images.

#### Background Correction

For every cycle and every channel, the background of the image was calculated by using a morphological opening operation with a 64x64 rectangular kernel. This background was subtracted from the respective channel before correcting for crosstalk and phasing effects.

### Basecalling

Input to basecalling is the detected beads $Beads = (B_1, ..., B_N)$ where $B_i \in R^{M\,x\,4}$ is bead $i$, $N$ is the number of detected beads and $M$ is the number of cycles. Every bead has a 4-dimensional intensity vector for every sequencing cycle.

#### Calculating the Crosstalk Matrix

For crosstalk correction, Optocoder uses the iterative correction method from[2]. Crosstalk Matrix $M$ is a 4x4 matrix which represents the interaction between channels. We calculate the crosstalk for all pairs of channels such that each corresponds to one element in the crosstalk matrix. This matrix is calculated from the bead intensities of the first cycle.

First, $M$ is initiated as an identity matrix. Then, the crosstalk for every pair of channels is calculated as below:

For a given pair of channels $(ch_a, ch_b)$:

1. The beads that have intensities between the 60th and 99th quantiles for the first component are chosen. $Beads^* = \{b_i \mid Q_A(0.60) < b_{i,1,a} < Q_A(0.99)\}$ where $A$ is the intensities of all beads in the first cycle for the first component channel and $b_{i,1,a}$ is the intensity of *channel a* of bead *i* at cycle 1.
2. Beads in $Beads^*$ are binned such that every bin includes ~10 beads.
3. A linear regression model is fitted to $Beads^*$ and the slope of this model is consecutively used to fill the crosstalk matrix entry for the respective pair.

The inverse of the crosstalk matrix is multiplied with the intensity data $Beads$ to calculate the corrected intensities for all beads. The whole process is repeated to update the crosstalk Matrix $M$, until either 15 iterations are reached or the maximum slope is lower than 0.05.

#### Generating the Phasing Matrix

Phasing matrix is defined with two values where $p$ is the probability or expected amount of phasing and $q$ is the expected amount of prephasing. Phasing matrix $P$ is initialised as an identity matrix of size (cycles x cycles) and is filled via a dynamic programming approach similar to[3].

$$P(i,j) = \begin{cases} p \cdot P(i-1,j) & j = 0 \\ p \cdot P(i-1,j) + (1-p-q) \cdot P(i-1,j-1) & j = 1 \\ p \cdot P(i-1,j) + (1-p-q) \cdot p(i-1,j-1) + q \cdot P(i-1,j-2) & \text{otherwise} \end{cases}$$

The values $p$ and $q$ can be selected by the user. Optocoder implements 0.07 phasing and 0.01 prephasing values by default.

### Basecalling

#### Crosstalk and Phasing Correction Model

The crosstalk and phasing correction model is defined as:

$$B_i = C S_i P,$$

where $B_i \in R^{M\,x\,4}$ is a matrix containing the observed intensities of bead $i$, $C \in R^{4\,x\,4}$ is the crosstalk matrix, $S_i \in R^{M\,x\,4}$ are the true intensities of bead $i$, and $P \in R^{M\,x\,M}$ is the phasing matrix, where $M$ is the number of cycles.

First, we calculate the crosstalk matrix and generate the phasing matrix as described above. Then, the inverse of the Kronecker product of the two matrices $(C \otimes P)^{-1}$ is calculated. This resulting correction matrix is multiplied with the bead intensities to acquire the corrected intensities.

$$S_i = (C \otimes P)^{-1} B_i \ . \qquad (1)$$

For the baseline basecallers, we use two different intensity profiles. For *naive basecalling,* we used the detected bead intensities after background correction without any crosstalk and phasing correction as:

$$S_i = B_i \ . \qquad (2)$$

For *crosstalk-only correction basecalling*, we first calculate the crosstalk matrix $C$ as described above and the phasing matrix $P$ is taken to be the Identity matrix.

$$S_i = (C \otimes I)^{-1} B_i \ . \qquad (3)$$

For the combined correction model, Phasing Matrix $P$ is computed as explained above with the default parameters and applied to (1).

#### Scaling and Basecalling

Before basecalling, we first scaled the intensity values for every channel using a Robust Scaler, defined as:

$$\frac{ch_{kl} - Q_1(ch_{kl})}{Q_3(ch_{kl}) - Q_1(ch_{kl})},$$

where $ch_{kl}$ are the intensities after corrections of all beads, $S$, for channel $k$ in cycle $l$, and $Q3$ and $Q1$ are the quartiles 3 and 1, respectively. Finally, we applied a SoftMax function to the final intensities for each bead. Finally, the bases are called for each bead and each cycle by taking the maximum intensity channel.

#### Phasing Parameter Search

To find the phasing and prephasing parameters that maximise the number of matches, we search for every pair of phasing and prephasing values between 0.0 and 1.0 with 0.01 step size. We calculated the number of matches for every pair and the parameter set with the maximum number of matches is selected. Then we run the whole pipeline again with the obtained optimised parameters.

### Quality Control Measures

#### Compression Score

Compression score for every barcode is calculated by computing the length of a compressed barcode. A barcode is compressed by computing the number of consecutive bases that are identical. For example, a repetitive, low complexity barcode such as "GGGGGGAAAAAA" would be compressed as "G6A6" while a complex barcode such as "GTACACATGCAC" would be compressed as "G1T1A1C1A1C1A1T1G1C1A1C1". In the QC plots, compression score distribution of a puck is compared against a theoretical distribution that is expected. The theoretical distribution is computed by calculating the compression score for a set of randomly generated barcodes of the same size with $BC$.

#### Shannon Entropy

The Shannon entropy for every barcode is calculated by:

$$H_{bc_i} = -\sum_{n \in bc_i} f(n, bc_i)^* \log_2(f(n, bc_i)),$$

where $f(n,\ bc_i)$ is the relative frequency of barcode $bc_i$ for $bc_i \in BC$. In the QC plots, entropy score distribution of a puck is compared against a theoretical distribution that is expected. The theoretical distribution is computed by calculating the Shannon entropy for a set of randomly generated barcodes of the same size with $BC$.

#### Chastity Score

We measure our base calling confidence by computing the chastity

$$C^{pq} = \frac{I^{pq}_{(n)}}{I^{pq}_{(n)} + I^{pq}_{(n-1)}} ,$$

where $I^{pq}_{(n)}$ and $I^{pq}_{(n-1)}$ are the intensities of the channels with the highest and the second highest values for bead $p$ in cycle $q$.

#### Registration Quality

Alignment score between two cycles is calculated by using the Structural Similarity Index (SSIM)[4] which measures the similarity between the reference image (i.e the last cycle) and the registered cycles by considering the texture information. This computation results in a value between 0-1 where 0 implies a completely different texture, 1 is achieved when there is a perfect match. In the QC plots, we show the scores for every cycle before and after registration where we choose 0.5 as an ad-hoc threshold to evaluate the registration quality of a run.

### In-house spatial transcriptomics method protocol

Samples S1-S4 were generated with an in-house spatial transcriptomics method that is based on Slide-Seq and Slide-SeqV2 protocols: an array with beads arranged on it is first generated, termed puck. The cellular barcodes are then optically decoded with a microscope as described in Slide-SeqV2[5]. ERCC RNA Spike-In Mix and a GFP reporter for S1 or E12 mouse brain sections for S2-S4 were subsequently placed on the optically decoded pucks and spatial transcriptomics libraries were generated following [5,6]. The prepared libraries were finally sequenced with NextSeq 500 Illumina machines and analysed as described below.

### Obtaining the Slide-Seq & Slide-SeqV2 optical sequencing barcodes

For Slide-Seq samples, we have downloaded the processed Puckcaller files from (https://singlecell.broadinstitute.org/single_cell/study/SCP354/slide-seq-study) and extracted the bead barcodes listed in the published AllBeadBarcodes.csv files for each sample. If there were multiple runs, we used the one with the latest date stamp. For the Slide-SeqV2 sample, we used the corresponding bead locations file from (https://singlecell.broadinstitute.org/single_cell/study/SCP815/highly-sensitive-spatial-transcriptomics-at-near-cellular-resolution-with-slide-seqv2).

### Processing & analysis of spatial transcriptomics sequencing data

To obtain the set of true barcodes of the samples S1-S4 we processed and analyzed the raw fastq files with spacemake v.0.4.2 with default parameters[7]. The cell barcode was extracted from read1 as the reverse sequence of nucleotide positions 9-20, while the UMI was extracted from read1 from nucleotide positions 1-8. Read2 was aligned to a combined genome of ERCC RNA Spike-In Mix and GFP reporter for S1 and to the mm10 genome for S2-S4. To obtain the set of true barcodes of the Slide-Seq and Slide-SeqV2 samples we used the tool DigitalGeneExpression from Drop-seq tools v.2.5.0 with default parameters to generate the digital gene expression matrix for the top 100,000 cell barcodes.

### Comparing Illumina-sequenced and the optically sequenced barcodes

After the Digital Gene Expression matrices were generated, we selected the top 100,000 bead barcodes with the highest number of counts to compare their nucleotide sequences against the sequences obtained by processing the optical sequencing data. For samples S1-S4 and Slide-SeqV2 samples, we counted exact matches between the two sets of barcodes. For Slide-Seq samples, images are generated through SOLiD sequencing and therefore they do not directly map to the nucleotides outputted by Optocoder. We have included a custom script that can convert the Illumina barcodes to the colour space by using the correct image and ligation sequences. Essentially, we ran Optocoder on all 20 images and took 14 non-constant cycles that are part of the barcode. While Optocoder outputs barcodes with nucleotides, we converted these barcodes to the numerical indices to represent them in the colour space. Then, the Illumina barcodes are converted to the colour space using the ligation sequences and orders as defined in https://github.com/MacoskoLab/PuckCaller and [6]. Finally, we counted exact matches between the two sets in colour space.

### Machine Learning Basecaller

#### Dataset

Machine learning basecaller is trained for every dataset separately provided that the respective Illumina sequencing run is available. Optocoder first calculates the matching barcodes between the optically decoded barcodes (after phasing correction) and the top 100,000 barcodes from the sequencing data. Specifically, input features for a sample is

$Beads^{Matching} = \{(B^s_i, bc_i) \mid \forall bc_i \in SB\}$ where $B^s_i$ is intensity matrix of bead *i* after robust scaling, $bc_i$ is the barcode corresponding to bead *i*, and $SB$ is the set of top 100,000 Illumina sequenced barcodes. Input dataset $Beads^{Matching}$ is used for training and is split into training (80%) and validation (20%) sets, and the non-matching barcodes are used as the test set.

### Models and Training

Optocoder implements four different types of classifiers. Random Forest (RF) and Multilayer Perceptron (MLP) models are implemented using scikit-learn (v0.22.2.post1), Gradient Boosting (GB) is implemented using XGBoost (v1.1.1), and finally a Recurrent Neural Network (RNN) model is implemented using Keras (v2.3.1) and Tensorflow (v2.1.0).

For RF, MLP and GB models, Optocoder uses scikit-learn's Multi Output Classifier approach where a classifier for each target is fitted. Here, one classifier for each sequencing cycle was trained. A random search was done with 10 iterations to optimise hyperparameters. Parameter ranges for the search are defined in the Optocoder machine learning module and the parameters of the selected models for the data presented in this paper are listed in Sup. Table 1. For the RNN model, we have one bidirectional LSTM layer, followed by dropout and a dense layer with softmax activation function. The model is trained using Adam optimizer and sparse categorical entropy loss. For every fit, 30 epochs with batch size 50 is used along with an early stopping mechanism. All samples were trained with these parameters. Furthermore, KerasTuner (v1.0.1) is used for the random hyperparameter search with 10 iterations. After training the models, new barcodes are predicted for non-matching beads. Optocoder saves the barcodes for all four models on disk, but the main basecaller is Gradient Boosting, as it consistently outperforms the others. Optionally, Optocoder machine learning module can be run with k-fold cross validation for parameter search and training steps.

### Barcode Prediction and Matching

After training the models, new barcode sequences are predicted for all optically decoded bead barcodes that do not match to any of the top 100,000 Illumina sequenced barcodes. The predicted barcodes that match to a sequenced barcode are then retained and added to the set of already matching barcodes.
