## Supplementary Tables for "Optocoder: computational decoding of spatially indexed bead arrays"

**Sup. Table 1. Machine Learning Model Parameters for the trained models**

| | P1 | P2 | P3 | P4 |
|---|---|---|---|---|
| XGB Classifier | **colsample_bytree** 0.795, **learning_rate** 0.236, **max_depth** 3, **min_child_weight** 4, **n_estimators** 911, **subsample** 0.849, **tree_method** 'hist' | **colsample_bytree** 0.767, **learning_rate** 0.517, **max_depth** 5, **min_child_weight** 3, **n_estimators** 477, **subsample** 0.781, **tree_method** 'hist' | **colsample_bytree** 0.817, **learning_rate** 0.070, **max_depth** 4, **min_child_weight** 4, **n_estimators** 489, **subsample** 0.396, **tree_method** 'hist' | **colsample_bytree** 0.855, **learning_rate** 0.213, **max_depth** 6, **min_child_weight** 1, **n_estimators** 550, **subsample** 0.734, **tree_method** 'hist' |
| Random Forest Classifier | **bootstrap** False, **max_depth** 110, **max_features** 'auto', **min_samples_leaf** 1, **min_samples_split** 2, **n_estimators** 180 | **bootstrap** False, **max_depth** 70, **max_features** 'auto', **min_samples_leaf** 2, **min_samples_split** 2, **n_estimators** 500 | **bootstrap** False, **max_depth** 50, **max_features** 'auto', **min_samples_leaf** 1, **min_samples_split** 5, **n_estimators** 180 | **bootstrap** False, **max_depth** 90, **max_features** 'sqrt', **min_samples_leaf** 1, **min_samples_split** 10, **n_estimators** 1000 |
| MLP Classifier | **hidden_layer_sizes** (100,), **activation** 'tanh', **solver** 'adam', **alpha** 0.05, **learning_rate** 'constant', **max_iter** 200 | **hidden_layer_sizes** (100,), **activation** 'tanh', **solver** 'adam', **alpha** 0.1, **learning_rate** 'adaptive', **max_iter** 200 | **hidden_layer_sizes** (50, 100, 50), **activation** 'tanh', **solver** 'sgd', **alpha** 0.1, **learning_rate** 'adaptive', **max_iter** 1000 | **hidden_layer_sizes** (100,), **activation** 'relu', **solver** 'adam', **alpha** 0.05, **learning_rate** 'constant', **max_iter** 1000 |
| RNN Classifier | **units**: 64 **dropout:** 0.1 **learning_rate:** 0.01 | **units:** 288 **dropout:** 0.2 **learning_rate:** 0.01 | **units:** 448 **dropout:** 0.3 **learning_rate:** 0.001 | **units:** 256 **dropout:** 0.2 **learning_rate:** 0.01 |

| | SSP1 | SSP2 | SSP3 | SSP4 |
|---|---|---|---|---|
| XGBClassifier | **colsample_bytree**<br>0.639,<br>**learning_rate**<br>0.410,<br>**max_depth**<br>3,<br>**min_child_weight**<br>1,<br>**n_estimators**<br>742,<br>**subsample**<br>0.732,<br>**tree_method**<br>'hist' | **colsample_bytree**<br>0.822,<br>**learning_rate**<br>0.278,<br>**max_depth**<br>5,<br>**min_child_weight**<br>1,<br>**n_estimators**<br>739,<br>**subsample**<br>0.726,<br>**tree_method**<br>'hist' | **colsample_bytree**<br>0.832,<br>**learning_rate**<br>0.316,<br>**max_depth**<br>4,<br>**min_child_weight**<br>4,<br>**n_estimators**<br>735,<br>**subsample**<br>0.825,<br>**tree_method**<br>'hist' | **colsample_bytree**<br>0.593,<br>**learning_rate**<br>0.366,<br>**max_depth**<br>3,<br>**min_child_weight**<br>2,<br>**n_estimators**<br>754,<br>**subsample**<br>0.772,<br>**tree_method**<br>'hist' |
| RandomForestClassifier | **bootstrap**<br>False,<br>**max_depth**<br>70,<br>**max_features**<br>'sqrt',<br>**min_samples_leaf**<br>2,<br>**min_samples_split**<br>2,<br>**n_estimators**<br>1000 | **bootstrap**<br>False,<br>**max_depth**<br>60,<br>**max_features**<br>'sqrt',<br>**min_samples_leaf**<br>1,<br>**min_samples_split**<br>5,<br>**n_estimators**<br>130 | **bootstrap**<br>False,<br>**max_depth**<br>100,<br>**max_features**<br>'sqrt',<br>**min_samples_leaf**<br>1,<br>**min_samples_split**<br>2,<br>**n_estimators**<br>500 | **bootstrap**<br>False,<br>**max_depth**<br>60,<br>**max_features**<br>'sqrt',<br>**min_samples_leaf**<br>1,<br>**min_samples_split**<br>2,<br>**n_estimators**<br>500 |
| MLPClassifier | **hidden_layer_sizes**<br>(50, 100, 50),<br>**activation**<br>'tanh',<br>**solver**<br>'sgd',<br>**alpha**<br>0.1,<br>**learning_rate**<br>'constant',<br>**max_iter**<br>600 | **hidden_layer_sizes**<br>(100,),<br>**activation**<br>'tanh',<br>**solver**<br>'adam',<br>**alpha**<br>0.0001,<br>**learning_rate**<br>'adaptive',<br>**max_iter**<br>600 | **hidden_layer_sizes**<br>(50, 100, 50),<br>**activation**<br>tanh',<br>**solver**<br>'sgd',<br>**alpha**<br>0.05,<br>**learning_rate**<br>'adaptive',<br>**max_iter**<br>400 | **hidden_layer_sizes**<br>(100,),<br>**activation**<br>'tanh',<br>**solver**<br>‘adam',<br>**alpha**<br>0.0001,<br>**learning_rate**<br>‘constant',<br>**max_iter**<br>200 |
| RNN Classifier | **units:** 288<br>**dropout:** 0.3<br>**learning_rate:** 0.001 | **units:** 448<br>**dropout:** 0.3<br>**learning_rate**:<br>0.01 | **units:** 448<br>**dropout:** 0.1<br>**learning_rate:**<br>0.01 | **units:** 352<br>**dropout:** 0.4<br>**learning_rate:**<br>0.001 |

**Sup. Table 2. Slide-Seq and Slide-Seq Data**

| Puck Name (Optocoder) | Puck ID | Slide-Seq Version |
|---|---|---|
| SSP1 | 180413_7 | V1 (SOLiD) |
| SSP2 | 180430_1 | V1 (SOLiD) |
| SSP3 | 180528_23 | V1 (SOLiD) |
| SSP4 | 200115_08 | V2 |
